## Supplemental Figure 1-2 for "Extracellular heparan 6-O-endosulfatases SULF1 and SULF2 in HNSC and other malignancies"

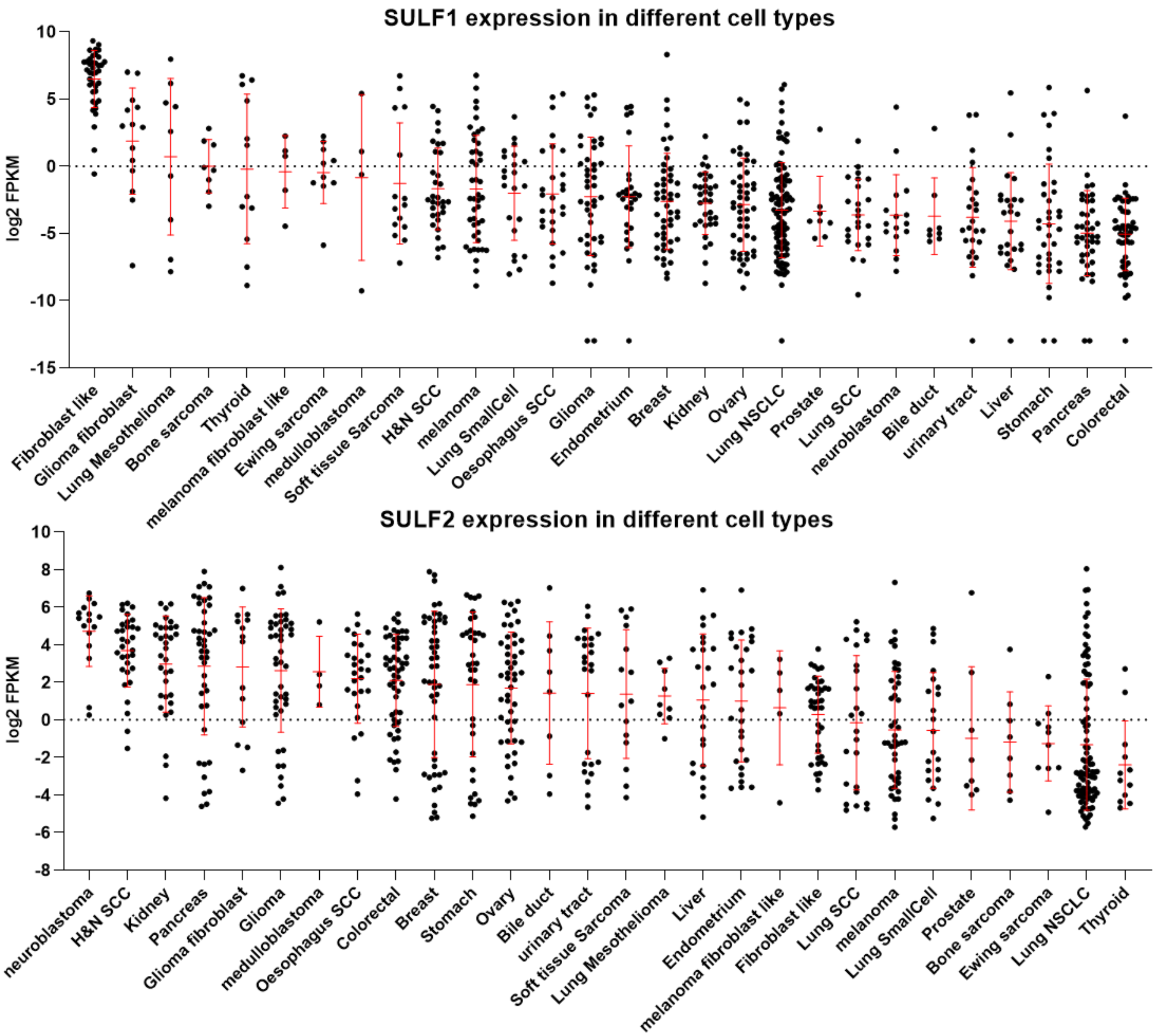


**Supplementary Figure 1.** A. SULF1 and B. SULF2 mRNA expression in cancer cell lines from the cancer cell-line encyclopedia (CCLE).

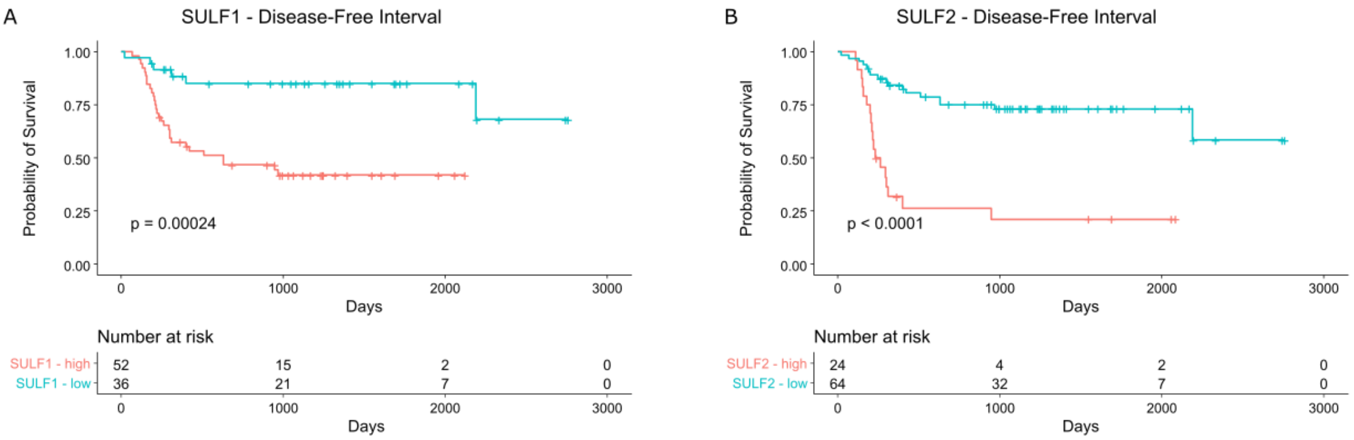


**Supplementary Figure 2.** Disease free survival of HNSC patients (n=88) based on the following: A. SULF1 expression; B. SULF2 expression.
